# Reinforcement Learning Is Impaired in the Sub-acute Post-stroke Period

**DOI:** 10.1101/2023.01.25.525408

**Authors:** Meret Branscheidt, Alkis M. Hadjiosif, Manuel A. Anaya, Jennifer Keller, Mario Widmer, Keith D. Runnalls, Andreas R Luft, Amy J. Bastian, John W. Krakauer, Pablo A. Celnik

## Abstract

**Background:** Neurorehabilitation approaches are frequently predicated on motor learning principles. However, much is left to be understood of how different kinds of motor learning are affected by stroke causing hemiparesis. Here we asked if two kinds of motor learning often employed in rehabilitation, (1) reinforcement learning and (2) error-based adaptation, are altered at different times after stroke.

**Methods:** In a cross-sectional design, we compared learning in two groups of patients with stroke, matched for their baseline motor execution deficit on the paretic side. The early group was tested within 3 months following stroke (N = 35) and the late group was tested more than 6 months after stroke (N = 30). Two types of task were studied: one based on reinforcement learning and the other on error-based learning.

**Results:** We found that reinforcement learning was impaired in the early but not the late group, whereas error-based learning was unaffected compared to controls. These findings could not be attributed to differences in baseline execution, cognitive impairment, gender, age, or lesion volume and location.

**Conclusions:** The presence of a specific impairment in reinforcement learning in the first 3 months after stroke has important implications for rehabilitation. It might be necessary to either increase the amount of reinforcement feedback given early or even delay onset of certain forms of rehabilitation training, e.g., like constraint-induced movement therapy, and instead emphasize others forms of motor learning in this early time period. A deeper understanding of stroke-related changes in motor learning capacity has the potential to facilitate the development of new, more precise treatment interventions.

## Introduction

Most upper limb motor recovery in humans and non-human animal models takes place early after stroke (∼1-3 months in humans, ∼1-3 weeks in rodents), a phenomenon that has been termed ‘spontaneous biological recovery’. Within this time window, responsiveness to training seems to be greater than outside of it in both animal models and in humans.^1,2^ Although the rapid recovery phenomenon occurs in most stroke survivors, its underlying physiological mechanisms in humans remain poorly understood.^3^

Animal models suggest that certain stroke-induced plasticity changes in cortex overlap with those seen during motor skill learning.^4,5^ For instance, different groups have described an enhancement of long term potentiation (LTP) in the peri-infarct tissue early after stroke.^6-8^ Similarly, motor skill learning and memory formation is dependent on LTP-mediated strengthening of synapses.^9-11^ The scarcity of motor learning studies at the acute/subacute stage after stroke is attributable in part to the performance confound - without matching for ability in task execution, differences in task performance can be misinterpreted as learning deficits.^3^ Thus, only careful matching for execution parameters allows for reliable conclusions about learning differences. For example, a recent study from Baguma and colleagues assessed motor skill learning in subacute stroke patients by changes in speed/accuracy trade-off in a tracking circuit task.^12^ Although the authors found that patients could learn the task, this study did not have a control group and did not control for differences in task execution, which may have under-estimated their learning ability and missed enhancement.

Given these observations, it remains an open question what to expect regarding motor learning being enhanced or not during the period of spontaneous biological recovery. Here, we investigated whether patients with stroke experience a higher capacity to learn movement via reinforcement (basal-ganglia associated) versus error-based mechanisms (cerebellar-associated), during and after the spontaneous biological recovery period. These two forms of learning were chosen because they are highly relevant in rehabilitation training.^13,14^ Critically, to assess learning we used a cross-sectional design to carefully match at baseline the ability to execute the tasks across groups. This would not be possible if we had studied the same patients longitudinally, as normal recovery would change their impairment level and they would no longer be naïve to the tasks.

## Methods

### Participants

We recruited 70 participants either within the first two months after stroke (*early group*), or ≥ six months after stroke (*late group*) and 17 age-matched healthy control participants from two centres (Johns Hopkins University, USA and Cereneo Centre for Neurology and Rehabilitation, CH). All patients met the following inclusion criteria: First-ever ischemic stroke with motor symptoms confirmed by imaging, supratentorial lesion location, one-sided upper extremity weakness (MRC < 5).

We excluded patients with minimal motor deficits in the first evaluation, defined as Fugl-Meyer Score of the upper extremity (FMS) >63/66 at recruitment (at the time point of testing two participants had recovered to FMS of 64 and one participants to a FMS of 65), age <21 years, hemorrhagic stroke or space-occupying hemorrhagic transformation, global inattention, visual field cut > quadrantanopia, receptive aphasia, inability to give informed consent or understand the tasks, other neurological or psychiatric illness that could confound execution/recovery. See Table 1 for details of patient characteristics.

**Table 1.**
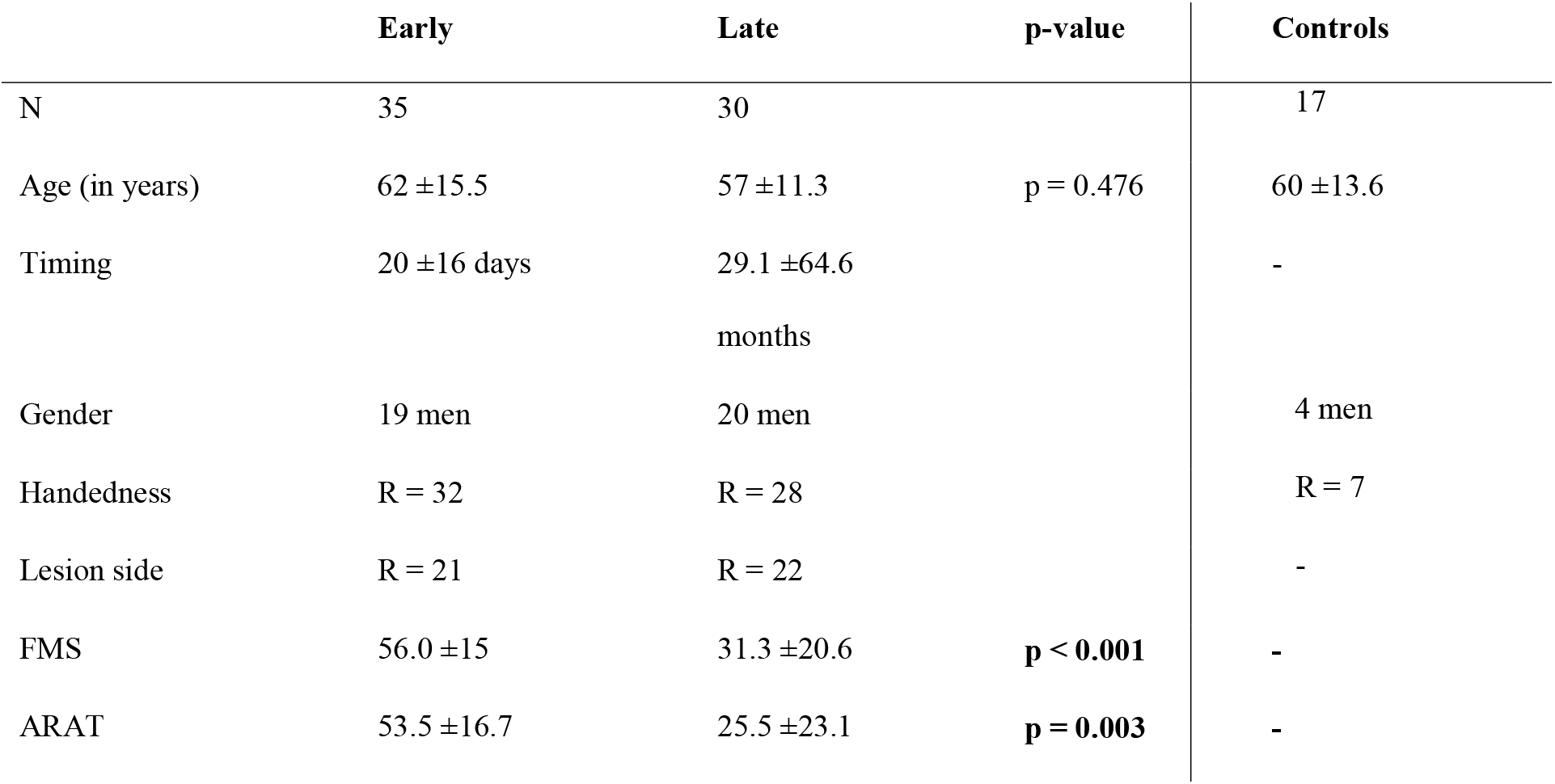
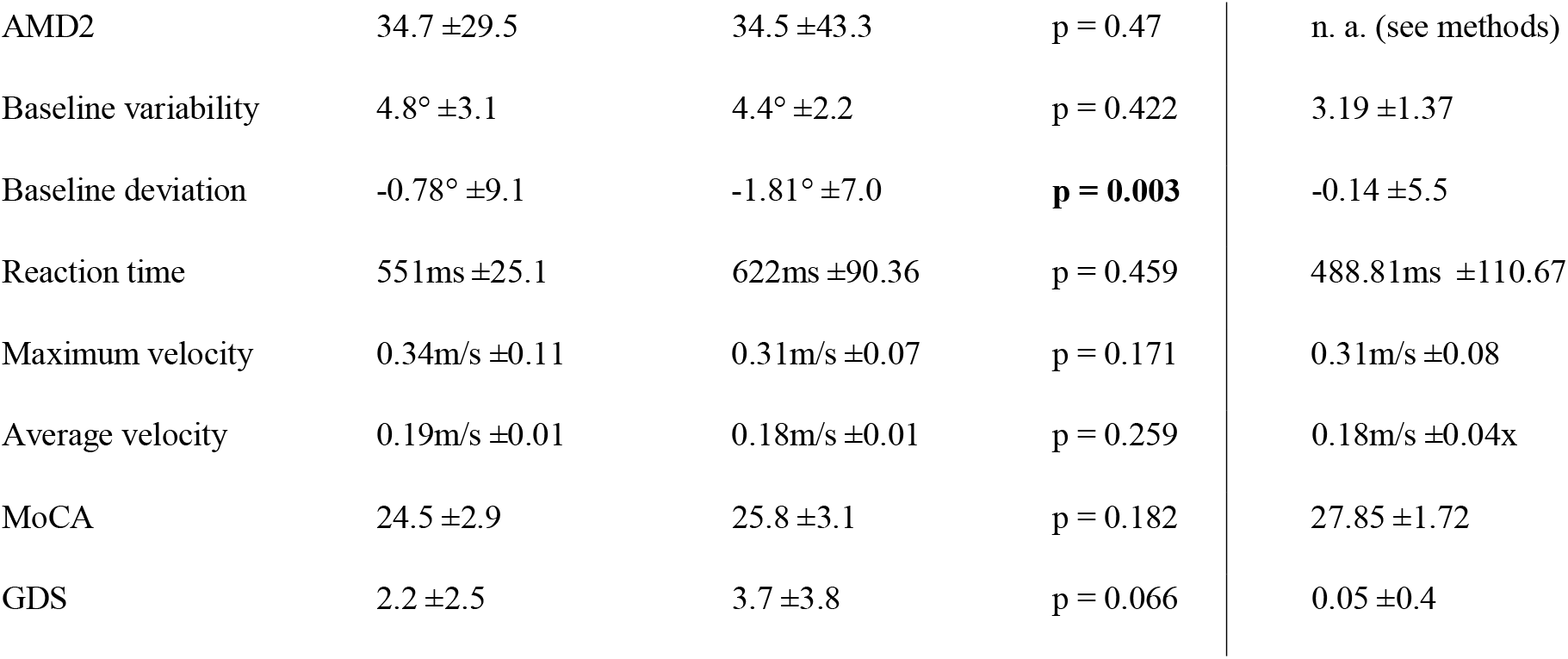
Clinical characteristics and reaching parameters. Median ±indicates standard deviation across participants. Timing: Timing of first assessment after stroke. FMS: Fugl-Meyer Score for the Upper Extremity; ARAT: Action Research Arm Test; AMD2: measurement of motor control (see methods); MoCA: Montreal Cognitive assessment; GDS: Geriatric Depression Scale

|  | **Early** | **Late** | **p-value** | **Controls** |
| --- | --- | --- | --- | --- |
| N | 35 | 30 |  | 17 |
| Age (in years) | 62 ±15.5 | 57 ±11.3 | p = 0.476 | 60 ±13.6 |
| Timing | 20 ±16 days | 29.1 ±64.6<br>months | | - |
| Gender | 19 men | 20 men |  | 4 men |
| Handedness | R = 32 | R = 28 |  | R = 7 |
| Lesion side | R = 21 | R = 22 |  | - |
| FMS | 56.0 ±15 | 31.3 ±20.6 | **p < 0.001** | - |
| ARAT | 53.5 ±16.7 | 25.5 ±23.1 | **p = 0.003** | - |
| AMD2 | 34.7 ±29.5 | 34.5 ±43.3 | p = 0.47 | n. a. (see methods) |
| Baseline variability | 4.8° ±3.1 | 4.4° ±2.2 | p = 0.422 | 3.19 ±1.37 |
| Baseline deviation | -0.78° ±9.1 | -1.81° ±7.0 | **p = 0.003** | -0.14 ±5.5 |
| Reaction time | 551ms ±25.1 | 622ms ±90.36 | p = 0.459 | 488.81ms ±110.67 |
| Maximum velocity | 0.34m/s ±0.11 | 0.31m/s ±0.07 | p = 0.171 | 0.31m/s ±0.08 |
| Average velocity | 0.19m/s ±0.01 | 0.18m/s ±0.01 | p = 0.259 | 0.18m/s ±0.04x |
| MoCA | 24.5 ±2.9 | 25.8 ±3.1 | p = 0.182 | 27.85 ±1.72 |
| GDS | 2.2 ±2.5 | 3.7 ±3.8 | p = 0.066 | 0.05 ±0.4 |

The experiments were approved by the ethics boards at Johns Hopkins School of Medicine Institutional Review Board and the Ethics Committee of Northwest and Central Switzerland in accordance to the Declaration of Helsinki and written informed consent was obtained from all participants.

### Study design

We chose a cross-sectional approach between early vs. late recovery periods to be able to match the participants’ ability to execute the motor tasks at baseline. This is critical in studies of motor learning in patients with stroke since it avoids differences in execution before training starts as well as the changes that might occur over time due to motor recovery. Within each group we tested outcome metrics at two time points one month apart. This was done to determine if rapid recovery changes that can be observed over a few weeks’ period affected learning. At each time point, we collected clinical and motor task data within seven days for the first time point (T1) or within one day for the second time point (T2; see Figure 1). To determine the total magnitude of learning that can be expected by a population in the same age group as our individuals with stroke, we assessed 17 age-matched, healthy participants with the same two reaching tasks.

**Figure 1.**
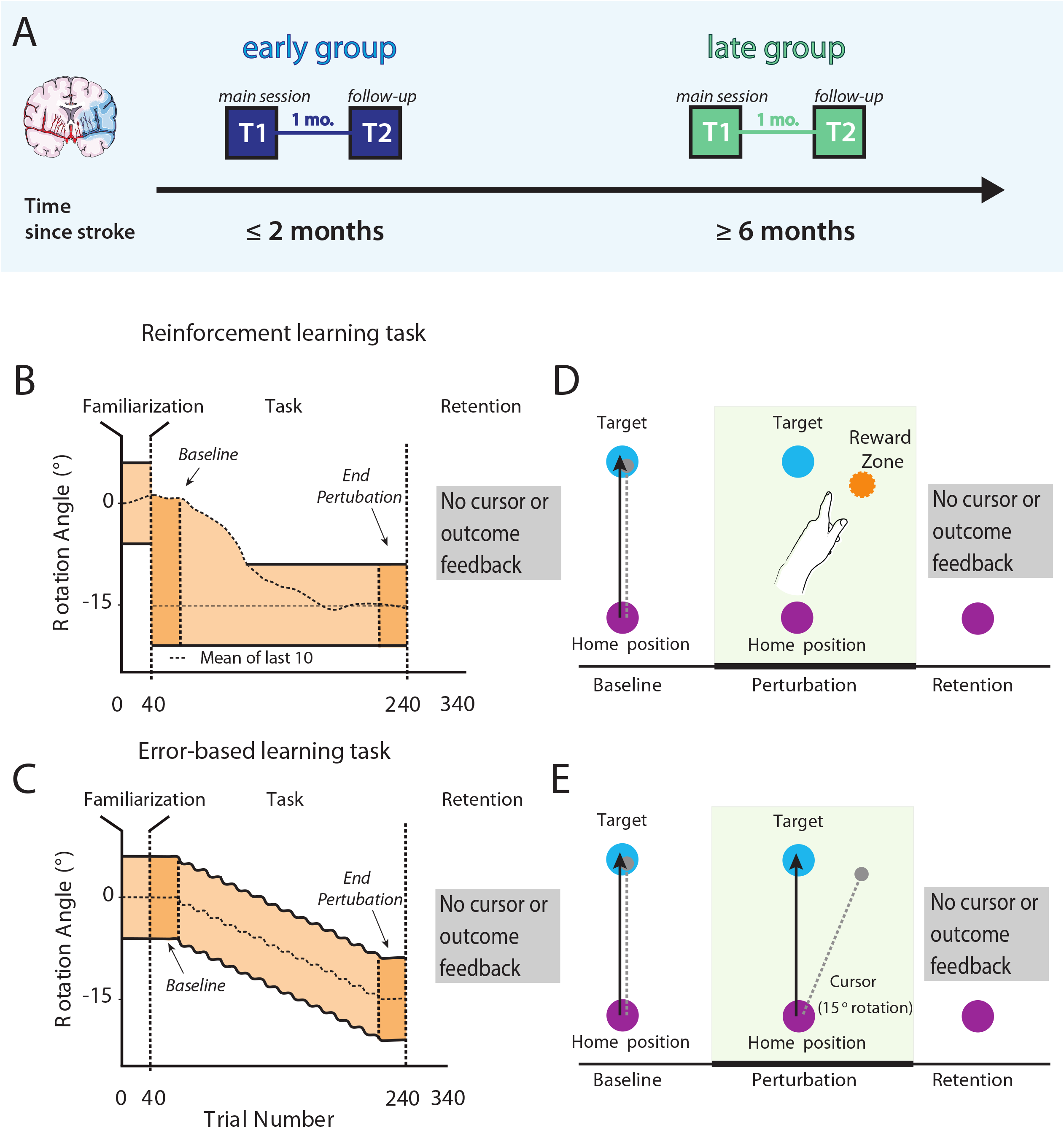
Study design and task overview with feedback conditions. (**A**) We recruited two separate groups of patients with stroke; one at the subacute stage (≤2 months, early group) and another one in the chronic period (≥6 months, late group). At the first time point (T1) participants were tested in two different motor learning tasks, a reinforcement-based motor task that relies predominantly on corticomotor-basal ganglia loops and a visuomotor error-based task that relies mostly on error-based learning processes driven by cerebellar plasticity. (**B & D**) In the reinforcement task no cursor feedback was provided, instead participants received only binary feedback about task success or failure if their reaches fell between the mean of the participant’s previous 10 reaches and the outer bound of the reward zone to -15°. (**C & E**) In the error-based learning task participants received online feedback on the cursor trajectory. After the baseline 40-trial period, a visuomotor rotation of 1 degree was imposed, and kept increasing by 2 degrees every 20 trials. Reward zone in both tasks is marked in light orange. To assess learning we compared *Baseline* and *End perturbation* trials (first and last 40 trials of task). After a baseline period to familiarize participants with the task, cursor rotation was gradually introduced until -15° in the error-based task. Within both groups clockwise or counter-clockwise rotation was counterbalanced, but later flipped for analysis. Image adapted after Therrien et al., 2016.

### Clinical data

The Fugl-Meyer Upper extremity score (FMS, max. score 66), and the Action Research Arm Test (ARAT, max. score 57), were used to assess impairment or functional deficits.^15,16^ Both measures were video recorded and graded by two trained assessors independently. Additionally, we collected the Montreal Cognitive Assessment (MoCA, max. score 30), and the Geriatric Depression scale (GDS, max. score 15), to capture cognitive impairment or depression, respectively.^17,18^

### Motor learning tasks

#### Set-up

To assess learning capacity, we investigated changes in performance in two different previously-published motor tasks; a *reinforcement* and an *error-based task*.^19,20^ Participants executed reaching movements with their paretic arm on a 2D plane while sitting in a KINARM exoskeleton robot (B-KIN Technologies) or the KINEREACH apparatus which provide antigravity support.^21^ We concealed arm movements by a screen and all visual feedback was projected on to the screen’s surface at the level of hand movement.

#### Evaluation of optimal reaching direction and measure of motor control

To ensure the ability to execute movements across groups was similar and prevent an execution confounder during learning, we matched patients’ motor control abilities at baseline using a global kinematic measure developed from a two-dimensional reaching task in a previous study.^22,23^ The task was designed to minimize the need for antigravity strength and prevent compensatory strategies.

Patients performed 10 cm point-to-point reaching movements from a home position to eight surrounding targets (176 reaches total/22 reaches per target in random order, target diameter 1cm, arrayed radially). The two adjacent target directions with the best execution (based on length of reach and successful target acquisition) were assigned as target directions for the motor learning tasks to optimize the ability to learn the tasks.

Using the same reaching data, we assessed motor control by using functional principal component analysis (fPCA) combined with the squared Mahalanobis distance (see Cortes et al.^23^ for a detailed description). This method computes a metric of the similarity between patients’ movement trajectories to those of a healthy, age-matched control group. The average squared Mahalanobis distance (AMD2) was then calculated for each individual and each target; we later used this metric to account for any execution confounder in our learning tasks.

#### Motor learning tasks

The reinforcement and error-based learning task used in this study have been adapted from Therrien et al.^19^ Instructions for both tasks were read to participants for standardization.

We instructed participants to make quick, 10-cm shooting movements from the home position through a single target. All participants performed a block to familiarize them with the task (40 trials). At this familiarization phase a white cursor represented the position of the index finger. To start a trial, the cursor had to be held stable in the start position light (purple, radius 1cm) before the target appeared (light blue circle, radius 1cm). The trials ended when the participant exceeded a distance of 10cm. To match movement times across both patient groups, we indicated too fast and too slow trials by a colour change of the target (<200ms = orange, >800ms = dark blue). Participants completed 340 reaches overall, over three blocks (Familiarization: 40 trials; Learning block: 40 Baseline and 160 Perturbation trials, Retention block: 100 trials).

In the perturbation phase, we introduced a gradual 15-degree rotation over 160 trials unbeknownst to the participant. Within both groups clockwise or counter clockwise rotations were counterbalanced. The rotation started after 40 trials. In both tasks, we did not provide any cursor or outcome feedback during the retention phase.

##### Reinforcement task

Here, participants did not get online cursor feedback, but only binary feedback for the trial outcome (the target turned green for successful hits and red for missed trials) based on the rotation angle. Outcome was based on comparing the reaching angle on the current trial with the moving average from the previous 10 trials. We provided reinforcement (green target) if the reaching angle was within the perturbed target direction (15° ± target width), or if the reaching angle was closer to the trained 15° perturbation compared to the moving average (Figure 1C). We provided failure signals (red target) if the opposite were true.

Because no online cursor feedback was available in these and all subsequent trials, the only information provided at the end of each trial was the successful acquisition of reinforcement (R+) or failure (R-). Trials that were too fast or too slow were given feedback unrelated to success/failure (the target turned light blue for trials that were too slow, and orange for trials that were too fast), discounted and instead repeated.

##### Error-based task

Here, participants received online feedback on the cursor trajectory. After the baseline 40-trial period, a visuomotor rotation of 1 degree was imposed, and kept increasing by 2 degrees every 20 trials.

Importantly, the magnitude of behavioral change expected after training (a 15° shift in reaching direction) was similar across tasks. Task order and perturbation direction were counterbalanced within groups.

### Data analysis

We flipped data from counter-clockwise sessions to analyze together with clockwise sessions. The number of trials that were repeated due to time violations in the reinforcement learning task was comparable in both groups.

We recorded hand position and velocity at the robotic handle at 1000 Hz (KINARM) or 420 Hz (KINEREACH) and analyzed offline with MATLAB. Following previous work, we chose to measure reach angle degree as the primary outcome metric. ^20,24-26^ We measured this from the start position to when the cursor crossed the target distance (10 cm away).

For both tasks we assessed learning in two different ways. First, we measured the difference between *Baseline* and *End perturbation* (average reaching angle for the last 40 trials of perturbation minus the first 40 trials of the baseline, *Total Learning*). Second, we assessed the total reaching angle deviation during *End perturbation* across groups.

Since motor learning includes two distinct processes, acquisition and retention, we assessed whether the time after stroke affected the magnitude and rate of retention across groups. To this end, we computed the difference in average reaching angle between the *End perturbation* and *Early Retention* trials (R1, first 40 trials of *Retention*) for the whole group and the baseline matched subgroup.

To rule out other execution factors that can result in learning differences across groups, we assessed *movement direction deviation* and *variability* at baseline (average reaching angle compared to 0° and average standard deviation of reaching angles at *Baseline*), *reaction time, average* and *maximum velocity* for all trials during the perturbation phase.

### Imaging

In a subset of patients that underwent clinical MRI, as a post-hoc analysis we evaluated lesion volume and location. A trained neurologist delineated manually lesion boundaries on each axial slice of a subject’s T2-weighted FLAIR or DWI image using MRICron software (http://www.mricro.com/mricron), see Figure 5 for averaged lesion distribution map. We normalized the obtained volume of interest (VOI) to the Montreal Neurological Institute (MNI) template using the clinical toolbox (http://www.nitrc.org/projects/clinicaltbx) with SPM12 (http://fil.ion.ucl.ac.uk/spm). We co-registered the T2 image to the T1 image and use these parameters to reslice the lesion into the native T1 space. We parcellated the brain in different regions (i.e., ROIs) using the JHU-MNI atlas.^27^ This atlas is implemented in the NiiStat software and contains 185 different ROIs covering the whole brain. To calculate the percentage of damage of the VOI for specific regions of interest (ROI) that have been implicated in reinforcement learning (orbitofrontal cortex, amygdala, caudate, putamen, nucleus accumbens and substantia nigra) we used NiiStat (www.niistat.org).

**Figure 2.**
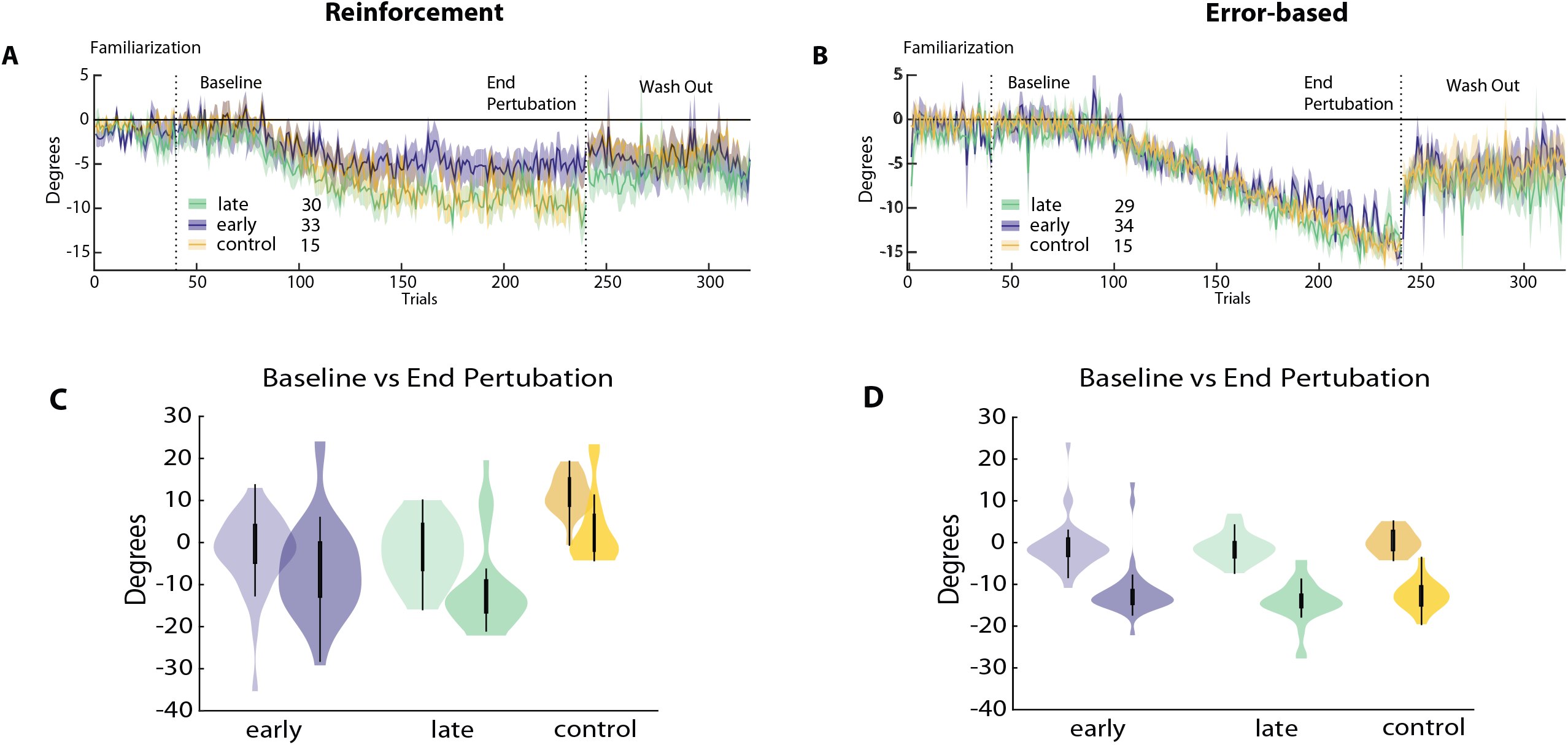
Reinforcement versus error-based learning at different stages after stroke. Results for the reinforcement task are on the left, error-based task on the right. (**A**) and (**B**) Changes in reaching angle over trials. (*Familiarization* = 40 trials, *Baseline* = 40 trials before introducing rotation, *End perturbation* = last 40 trials of rotation, *Wash out* = 100 trials without any feedback). Early group in blue, late group in green. (**C**) and (**D**) Comparison for *Baseline* versus *End perturbation* in both groups.

**Figure 3.**
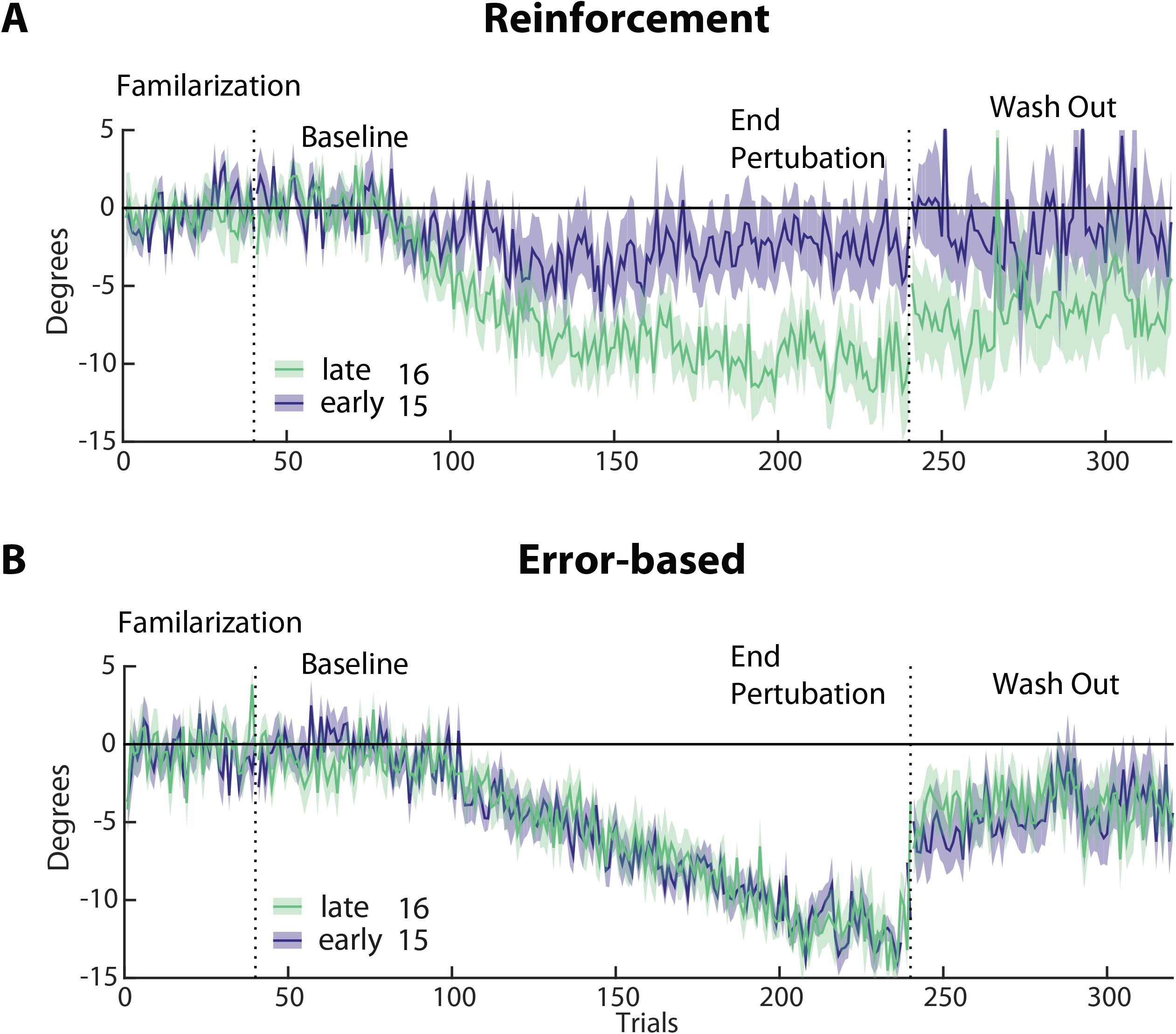
Subgroup analysis of reinforcement versus error-based learning in the early (blue, n=15) and late (green, n=16) groups. **(A & B)** Changes in reaching angle over trials. (*Familiarization* = 40 trials, *Baseline* = 40 trials before introducing rotation, *End perturbation =* last 40 trials of rotation, *Wash Out* = 100 trials without any feedback). Please note the reduced learning in the early vs. late group in the reinforcement task only. Shading indicates SEM.

**Figure 4.**
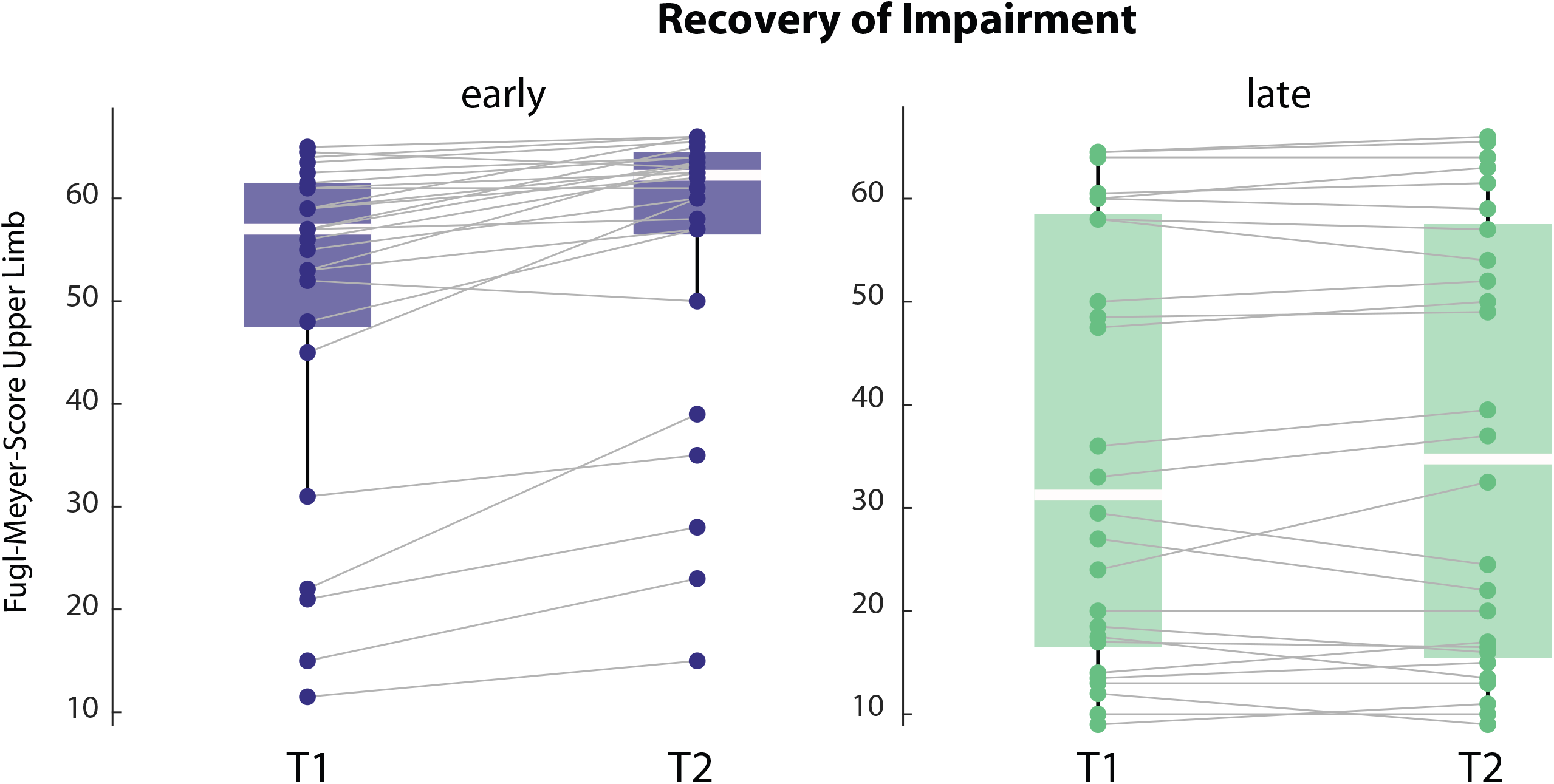
Recovery trajectory for impairment, measured by Fugl-Meyer Score for the upper limb at T1 and T2, in the early and the late groups. Note that early group improves over time and has overall lower impairment.

**Figure 5.**
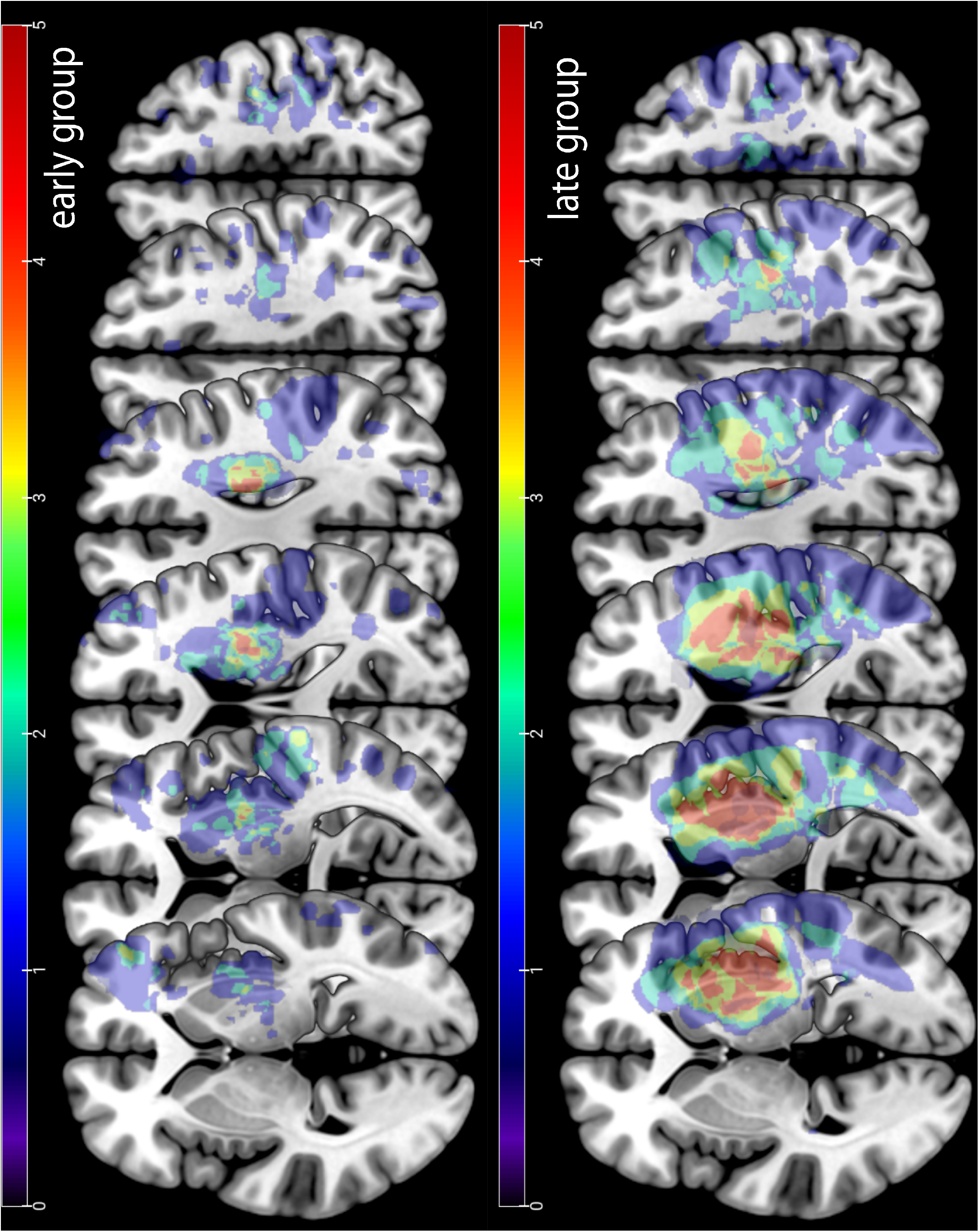
Stroke lesion overlay. Upper row lesion location for the early group. Lower row lesion location for the late group. Note that the late group had overall larger lesion volume than the early group.

### Statistics

Because the assumption for normality was not fulfilled for most reaching related variables, we use permutation testing to assess differences between groups. We reassigned participants randomly to either the ‘early’ or ‘late’ group, and the difference between the resampled groups was computed. This procedure was repeated 10,000 times, allowing us to generate a null distribution assuming no group differences. The proportion of resampled values that exceeded the true observed difference was used to compute p-values and determine statistical significance. Under the null hypothesis, the true difference between the two groups should lie within the distribution of these randomly generated differences, with extreme values providing evidence against the null hypothesis. We used this approach for all outcome variables unless explicitly stated differently. We used the same methods for the comparison of both stroke groups and the healthy control participants.

Total lesion volume as well as percentage of damage for the different ROI implicated in reinforcement learning was compared between groups using a simple two-sided Student’s t-test and, where appropriate, multiple testing correction was performed. All data are expressed as median ±standard deviation unless stated otherwise. Statistical analyses were performed using custom-written MATLAB and R routines.

### Data availability

Data and custom-written code will be available upon publication on an open repository.

## Results

A total of 70 patients were enrolled in the study. We excluded five participants (one person because of a MoCa score below 20, indicating significant cognitive impairment, two persons because of protocol time violation, and two because of technical problems). The early group included 35 participants (median 20 days after the insult, range 6 - 58 days), whereas the late group included 30 participants (median 29.1 months after stroke, range 7 months – 30 years). We also collected data from healthy control participants (N = 17; see Table 1 for clinical characteristics per group and Table 2 per individual participant).

**Table 2.**
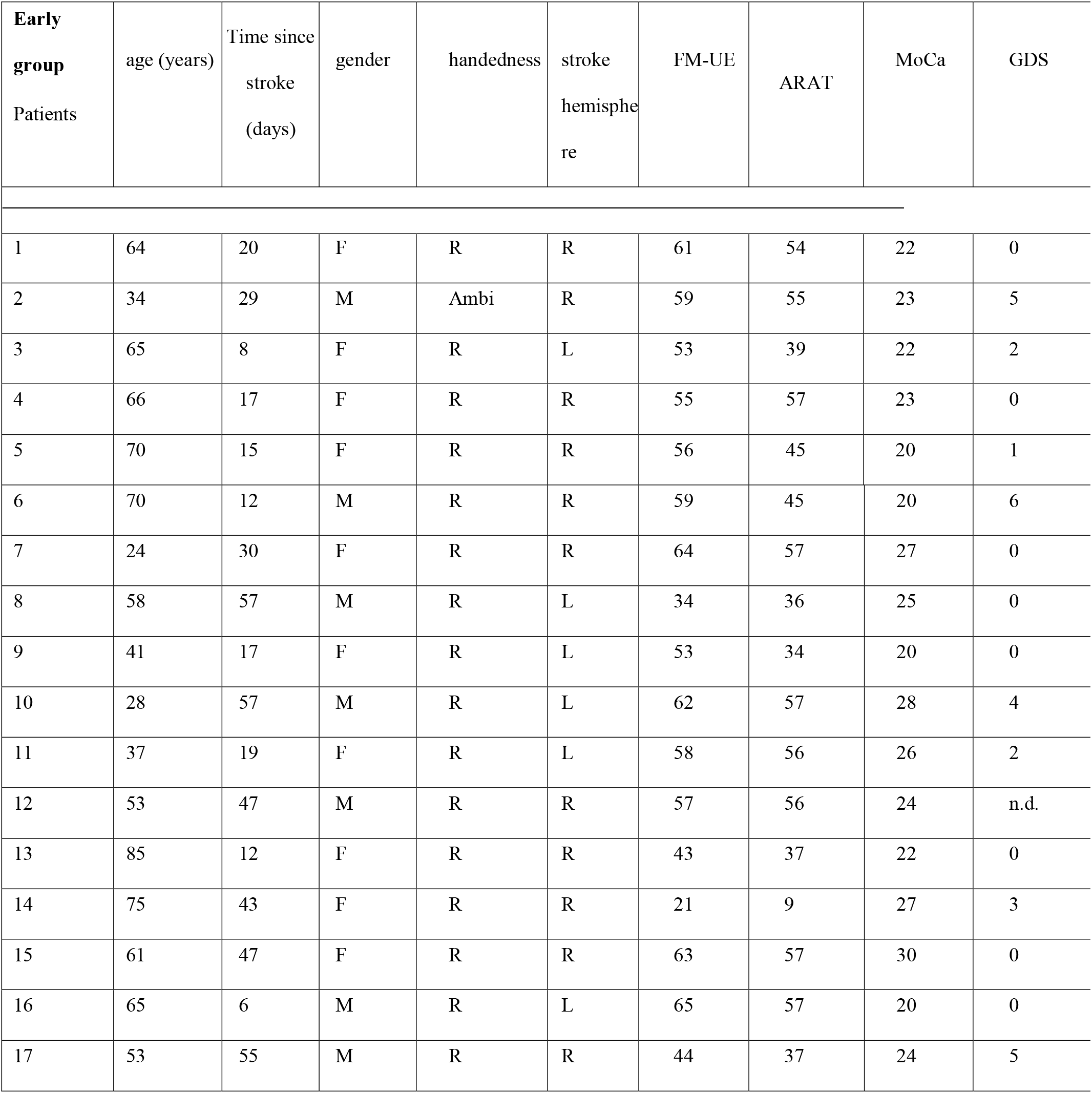
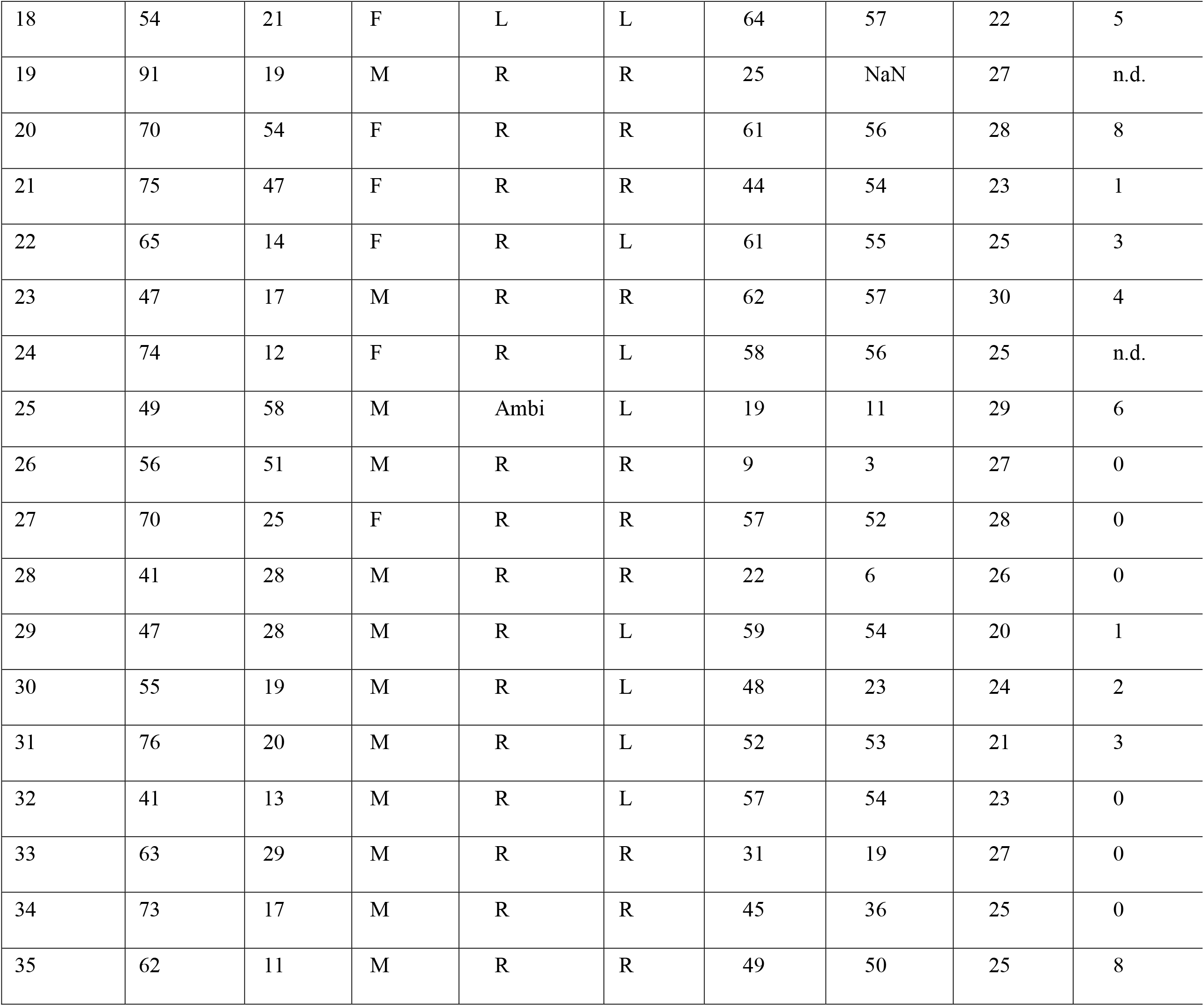
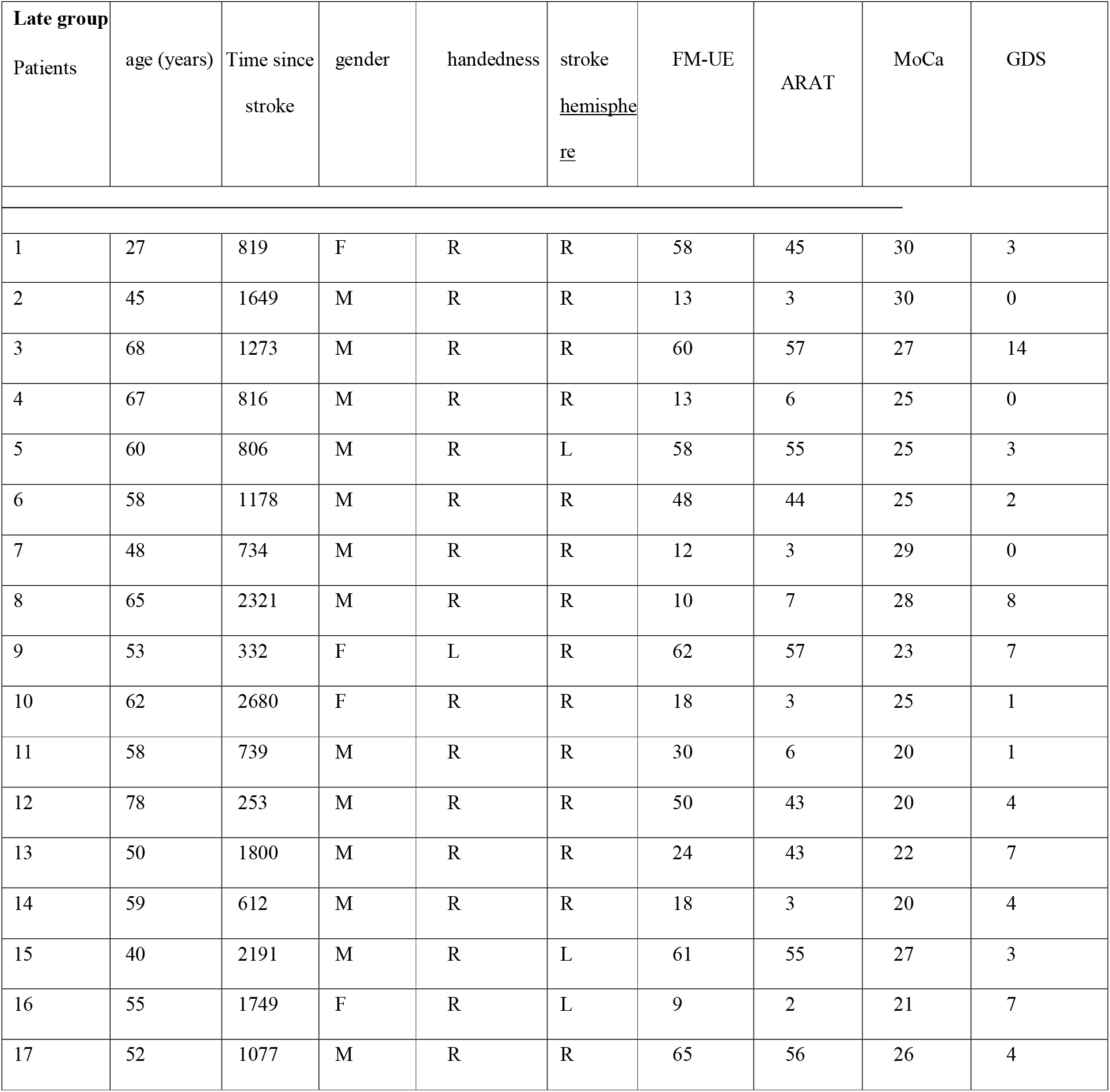
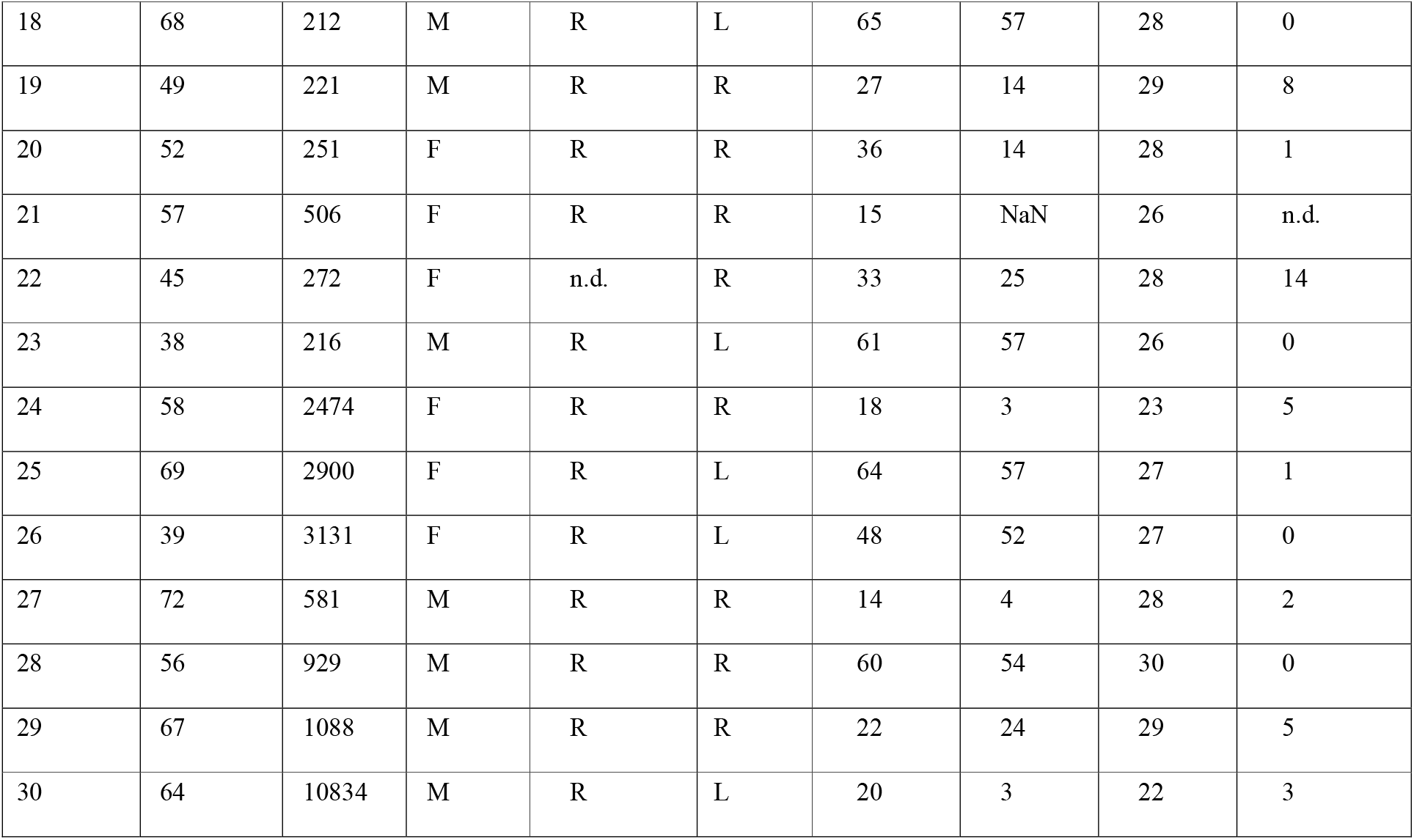
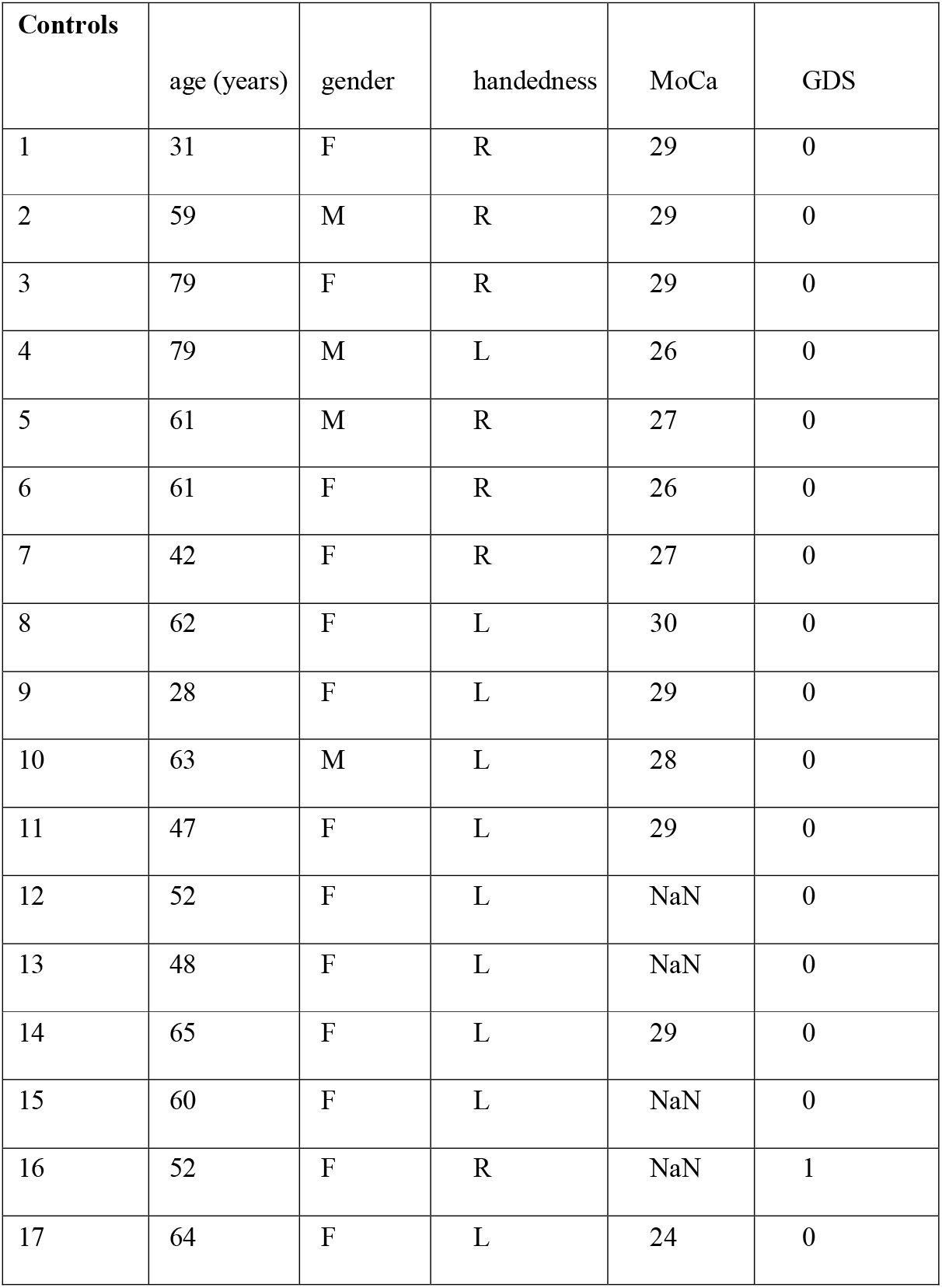
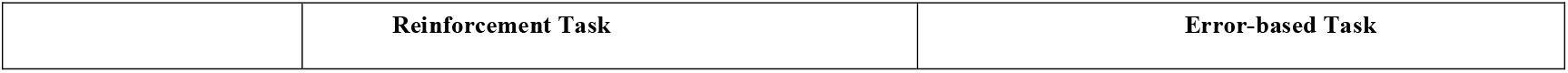
Patient and healthy control characteristics: age (years), time since stroke (days), gender, handedness, paretic side, initial FMS (Fugl-Meyer upper limb score, maximum 66), ARAT and initial MoCA (Montreal Cognitive Assessment, maximum 30).

Data collection problems occurred during the reinforcement task in two patients from the early group and in the error-based task for one person of the late group (note the exact N of participants for each task in Figure 2 and 3). For an overview of all results see also Table 3.

**Table 3.**
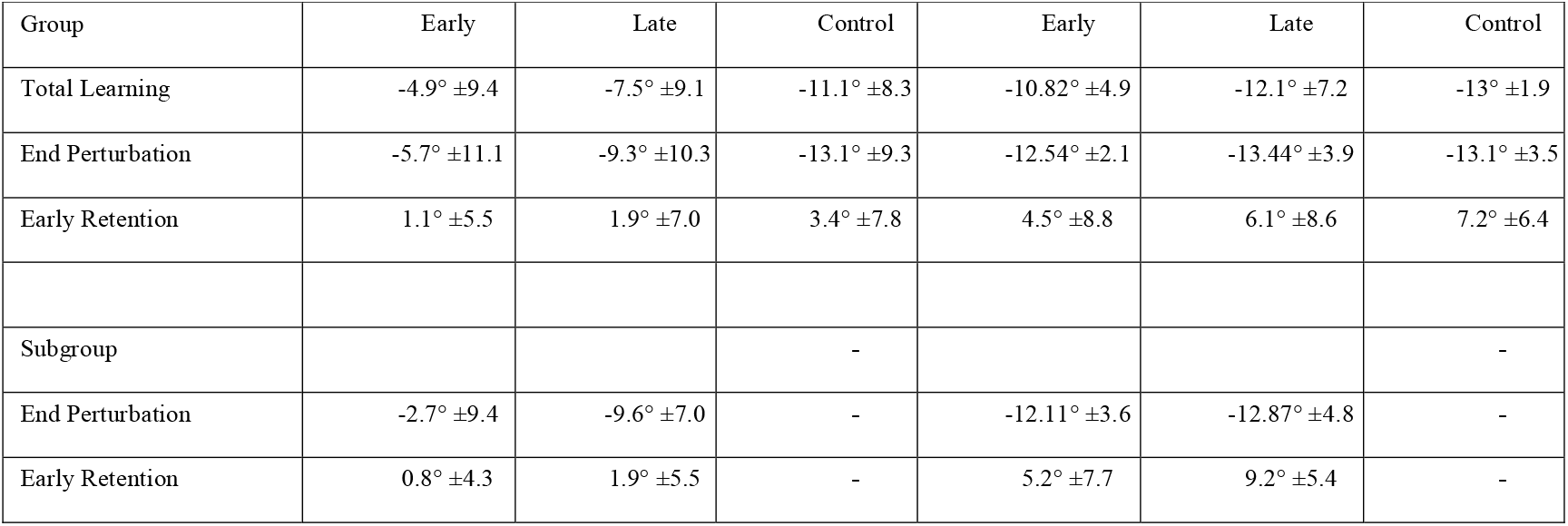
Outcomes per task of each group. Median ±indicates standard deviation across participants. *Total Learning*: average reaching angle for the last 40 trials of perturbation minus the first 40 trials of the baseline, *End Perturbation*: average reaching angle for the last 40 trials of perturbation, *Early Retention:* first 40 trials of the retention block. *Subgroup analysis:* participants from both stroke groups with matched *Baseline* performance.

### Reinforcement Motor learning capacity was reduced in the early group

Surprisingly, we found significantly less learning in the reinforcement task at the *End perturbation* in the early versus the late group and healthy controls (early: -5.7° ±11.1 versus late: -9.3° ±10.3, *p* = 0.035; early versus controls -13.1° ±9.3, *p* = 0.049). In addition, *Total Learning* was not higher, but in fact was less in the early versus the late group and controls (early: -4.9° ±9.4 versus late -7.5° ±9.1, *p* = 0.033; early versus controls -11.1° ±8.3, *p* = 0.048).

Despite patients being matched for motor control abilities at baseline (see Methods) and displaying no difference in other factors that could possibly affect task execution (see under Clinical and Kinematic variables cannot explain learning differences), the two patient groups differed significantly in their average reaching angles at *Baseline* (early: -0.78° ±9.1, late: -1.81° ±7.0, *p* = 0.003). This means that the *Total Learning* difference across groups could have been driven by this difference at baseline. Thus, to account for reaching angle execution at baseline, we conducted a subgroup analysis that included participants from both groups with matched *Baseline* execution (average reaching angle ±5°, resulting in *N* = 16 in the early group and N=15 in late group). This subgroup analysis confirmed that the early group had markedly less *Total Learning* compared to the late group (see Figure 3A; early: -2.7° ±9.4 versus late: -9.6° ±7.0, *p* = 0.03). Indeed, matching baseline execution also highlighted the difference in *End perturbation* angles between groups (early: -2.4° ±9.2 versus late: -9.9° ±8, *p* = 0.025).

### Error-based motor learning capacity after stroke was not different from healthy controls

As predicted, we found a comparable average shift in reach angles at *End perturbation* between groups in the *error-based* task (early: -12.54° ±2.1 versus late: -13.44° ±3.9, *p* = 0.18; early versus controls -13.1° ±3.5, *p* = 0.56). *Total Learning* in this task was similar between the early, the late and the control groups (early: -10.82° ±4.9 versus late: -12.1° ±7.2, *p* = 0.18; early versus control: -13° ±1.9, *p* = 0.21). Importantly, the *subgroup analysis* in the matched-baseline group also did not show statistical difference (early: -12.11° ±3.6 versus late: -12.87° ±4.8, *p* = 0.191; Figure 3B).

Of note, the percentage of trials that were repeated because of timing issues was under 10% and did not differ between groups (early: 7%, late 5%, t (50) = 1.21, *p* = 0.23).

### There were no post-stroke abnormalities in retention following training in the reinforcement and error-based tasks

For the reinforcement task, we found no significant difference in R1 across groups (early: 1.1° ±5.5 versus late: 1.9° ±7.0, *p* = 0.43), even when matched for baseline reaching angle deviation (early: 0.8° ±4.3 versus late: 1.9° ±5.5, *p* = 0.49). Compared to healthy controls, the early group had a significantly smaller R1, however this effect might be attributed to the smaller deviation in *End perturbation* in the first place (control: 3.4° ±7.8, *p* = 0.041).

We found similar results for the error-based task for the whole group (early: 4.5° ±8.8 versus late: 6.1° ±8.6, *p* = 0.23, early versus control: 7.2° ±6.4, *p* = 0.83) or the baseline-matched subgroup (early: 5.2° ±7.7 versus late: 9.2° ±5.4, *p* = 0.125).

### Group differences in clinical and kinematic variables could not explain the differential effect of stroke on reinforcement and error-based learning

To determine whether differences in learning capacity at the early versus late stage after stroke were due to other variables beyond learning capacity, we assessed factors that could possibly affect task execution. Despite all participants having similar abilities in executing the reaching tasks (AMD^2^, baseline variability and reaching execution parameters, see methods), early group had less overall motor impairment (measured by the FMS scores) and better functional deficits (measured by the ARAT) than those in the late group. Importantly, neither FMS (R = 0.09, p = 0.09) nor ARAT (R = - 0.02, p = 0.41) correlated with Total Learning. Finally, cognitive function and mood disturbances were similar across both groups (Table1).

### Motor recovery followed the expected longitudinal pattern

To ensure our participants followed the expected normal recovery pattern of rapid motor impairment changes early but not late (>6months) after stroke we compared FMS scores at the time of the motor learning testing (T1) and a second time point one month later (T2). In addition, this comparison can help understand whether the lower reinforcement capacity observed early after stroke impacted our participants’ recovery. We found that patients had a phenotypical change, where the early group showed a significant increase in FMS scores over time, while the late group remained stable (see Figure 4, of note only patients with both time points were included; early *N* = 23, late *N*= 28; *FMS*, early T1: 49.3 ±14.9, T2: 55.3 ±14.4, t = -5.324, p < 0.001; late T1: 36.3 ±20.6, T2: 36.1 ±20.7, t = -0.51, p = 0.614).

### Learning one month later still showed differences between both groups

We also assessed learning in the reinforcement task in both groups at T2. *Total Learning* and *End perturbation* one month later was still lower in the early compared to the late group. However, the differences were not statistically significant due to an improvement of the early group while the late group remained stable in their performance (*Total Learning*: early -6.9° ±7.6 versus late -8.2° ±7.8, *p* = 0.055; *End perturbation*: early - 7.7° ±10.8 versus late -8.5° ±9.0, *p* = 0.257). Importantly, since one third of the participants were lost to follow-up, this comparison across the whole group needs to be taken with caution.

### Imaging analysis

To test whether lesion location could explain the abnormal performances in the reinforcement task, we performed a post-hoc analysis on total lesion volume and lesion volume of regions of interest (ROI) known to be involved in reinforcement processing between the two groups. As this imaging analysis was not part of the original study protocol, MRI data was only available for a subset of participants; 25 participants in the early and 15 participants in the late group. Overall lesion volume was statistically larger in the late group compared to the early group (*p* = 0.018, Figure 5), a finding consistent with the higher motor impairment (worse FM scores) in the late group.

To determine the potential impact of damage in the regions of interest implicated in the neural circuitry underlying reinforcement learning, we assessed the percentage of lesion load in six ROIs: orbitofrontal cortex, amygdala, caudate nucleus, putamen, nucleus accumbens and substantia nigra.^28^ In each of these brain regions percentage of lesion load was higher in the late compared to the early group, making it very unlikely that unbalanced lesion distribution could account for the differences in reinforcement learning (early versus late group, orbitofrontal: *N* = 3 vs 3, amygdala: *N* = 1 vs 4, caudate nucleus *N* = 6 vs 8, putamen *N* = 10 vs 7, nucleus accumbens *N* = 0 vs 2 and substantia nigra *N* = 1 vs 2).

## Discussion

Here, we asked whether there are differences in motor learning ability at two time points after stroke. Specifically, we examined two types of motor learning during the subacute (<3months, early group) and chronic (>6months, late group) period. The first one was a reinforcement learning and the second one was a visuomotor error-based adaptation.

We found that in the sub-acute post-stroke period reinforcement motor learning capacity was lower compared to patients in the chronic phase. Error-based learning in contrast was comparable at both time points, indicating that the deficit in reinforcement learning was not merely evidence for a more general learning deficit. Importantly, the motor learning findings could not be explained by differences in baseline ability to execute the two tasks, as patients in the two groups were carefully matched for performance, nor differences in either overall disability or lesion volume. Of note, the capacity to learn via reinforcement in the late group (i.e. >6months following stroke) was similar to the healthy age-matched group. Based on these findings, it appears that the two forms of motor learning investigated here do not follow the time courses for either spontaneous biological recovery or responsiveness to rehabilitative training seen in animal models or more recently in humans.^1,2^

Our findings have important implications for the development of rehabilitation strategies following stroke. Dromerick and colleagues showed that patients responded better in the first three months than after six months.^2^ Our results suggest that it might not only be a matter of timing, but also of what learning mechanisms are in play. Thus, it may be better to deemphasize interventions based on reinforcement learning in the sub-acute stroke period. The results of the EXCITE trial may appear to contradict this, however the reported positive effects of early CIMT training after stroke are likely explained by higher intensity compared to usual care rather than therapy content. ^28^ Furthermore, they fall into a period (3 to 9 months post-stroke) that would blend the early and late group tested within our framework.

## Limitations

This study probed two forms of motor learning frequently used in motor rehabilitation of the upper extremity. The results do not contradict the animal literature showing higher sensitivity to training in a critical period after stroke.^29^ This is because many training protocols involve learning mechanisms that go beyond either reinforcement learning with exogenous reward or visuomotor adaptation. In addition, rehabilitation training extends for periods much longer than the time it takes to just assay a specific learning mechanism. Furthermore, we tested acquisition and retention processes, yet there are other phenomena that are associated with motor learning, such as consolidation and savings, that have not been explored here.

## Conclusion

The observation that reinforcement learning is impaired early after stroke has important implications for the design of novel rehabilitation interventions. Training protocols will need to consider a timing x learning mechanism interaction. For instance, rehabilitation exercises might need to weight more error-based, strategic or skill learning during the sub-acute period after stroke, while minimizing reliance on reinforcement and/or use this form of learning later in the recovery process. It is also possible that augmenting reinforcement signals could be necessary to compensate for this deficit.

## Acknowledgements

We would like to thank our patients willing to participate in this research.

## Funding

This work was supported by NIH grant 5R01HD053793.

## Potential conflict of interest

None of the authors have a conflict of interest to declare.

## Notes

### Competing Interest Statement

The authors have declared no competing interest.

