## Supplementary material for "Reinforcement Learning Is Impaired in the Sub-acute Post-stroke Period": Table 1

**Table 1 Clinical characteristics and reaching parameters**

|  | **Early** | **Late** | **p-value** | **Controls** |
| --- | --- | --- | --- | --- |
| N | 35 | 30 |  | 17 |
| Age (in years) | 62 ±15.5 | 57 ±11.3 | p = 0.476 | 60 ±13.6 |
| Timing | 20 ±16 days | 29.1 ±64.6 months |  | - |
| Gender | 19 men | 20 men |  | 4 men |
| Handedness | R = 32 | R = 28 |  | R = 7 |
| Lesion side | R = 21 | R = 22 |  | - |
| FMS | 56.0 ±15 | 31.3 ±20.6 | **p < 0.001** | **-** |
| ARAT | 53.5 ±16.7 | 25.5 ±23.1 | **p = 0.003** | **-** |
| AMD2 | 34.7 ±29.5 | 34.5 ±43.3 | p = 0.47 | n. a. (see methods) |
| Baseline variability | 4.8° ±3.1 | 4.4° ±2.2 | p = 0.422 | 3.19 ±1.37 |
| Baseline deviation | -0.78° ±9.1 | -1.81° ±7.0 | **p = 0.003** | -0.14 ±5.5 |
| Reaction time | 551ms ±25.1 | 622ms ±90.36 | p = 0.459 | 488.81ms ±110.67 |
| Maximum velocity | 0.34m/s ±0.11 | 0.31m/s ±0.07 | p = 0.171 | 0.31m/s ±0.08 |
| Average velocity | 0.19m/s ±0.01 | 0.18m/s ±0.01 | p = 0.259 | 0.18m/s ±0.04x |
| MoCA | 24.5 ±2.9 | 25.8 ±3.1 | p = 0.182 | 27.85 ±1.72 |
| GDS | 2.2 ±2.5 | 3.7 ±3.8 | p = 0.066 | 0.05 ±0.4 |

Median ±indicates standard deviation across participants. Timing: Timing of first assessment after stroke. FMS: Fugl-Meyer Score for the Upper Extremity; ARAT: Action Research Arm Test; AMD2: measurement of motor control (see methods); MoCA: Montreal Cognitive assessment; GDS: Geriatric Depression Scale
