## Supplementary material for "Reinforcement Learning Is Impaired in the Sub-acute Post-stroke Period": Table 2

**Table 2 Patient and healthy control characteristics**: age (years), time since stroke (days), gender, handedness, paretic side, initial FMS (Fugl-Meyer upper limb score, maximum 66), ARAT and initial MoCA (Montreal Cognitive Assessment, maximum 30).

| **Early group**  Patients | age (years) | Time since stroke (days) | gender | handedness | stroke hemisphere | FM-UE | ARAT | MoCa | | GDS |
| --- | --- | --- | --- | --- | --- | --- | --- | --- | --- | --- |
| 1 | 64 | 20 | F | R | R | 61 | 54 | 22 | | 0 |
| 2 | 34 | 29 | M | Ambi | R | 59 | 55 | 23 | | 5 |
| 3 | 65 | 8 | F | R | L | 53 | 39 | 22 | | 2 |
| 4 | 66 | 17 | F | R | R | 55 | 57 | 23 | | 0 |
| 5 | 70 | 15 | F | R | R | 56 | 45 | 20 | | 1 |
| 6 | 70 | 12 | M | R | R | 59 | 45 | | 20 | 6 |
| 7 | 24 | 30 | F | R | R | 64 | 57 | | 27 | 0 |
| 8 | 58 | 57 | M | R | L | 34 | 36 | | 25 | 0 |
| 9 | 41 | 17 | F | R | L | 53 | 34 | | 20 | 0 |
| 10 | 28 | 57 | M | R | L | 62 | 57 | | 28 | 4 |
| 11 | 37 | 19 | F | R | L | 58 | 56 | | 26 | 2 |
| 12 | 53 | 47 | M | R | R | 57 | 56 | | 24 | n.d. |
| 13 | 85 | 12 | F | R | R | 43 | 37 | | 22 | 0 |
| 14 | 75 | 43 | F | R | R | 21 | 9 | | 27 | 3 |
| 15 | 61 | 47 | F | R | R | 63 | 57 | | 30 | 0 |
| 16 | 65 | 6 | M | R | L | 65 | 57 | | 20 | 0 |
| 17 | 53 | 55 | M | R | R | 44 | 37 | | 24 | 5 |
| 18 | 54 | 21 | F | L | L | 64 | 57 | | 22 | 5 |
| 19 | 91 | 19 | M | R | R | 25 | NaN | | 27 | n.d. |
| 20 | 70 | 54 | F | R | R | 61 | 56 | | 28 | 8 |
| 21 | 75 | 47 | F | R | R | 44 | 54 | | 23 | 1 |
| 22 | 65 | 14 | F | R | L | 61 | 55 | | 25 | 3 |
| 23 | 47 | 17 | M | R | R | 62 | 57 | | 30 | 4 |
| 24 | 74 | 12 | F | R | L | 58 | 56 | | 25 | n.d. |
| 25 | 49 | 58 | M | Ambi | L | 19 | 11 | | 29 | 6 |
| 26 | 56 | 51 | M | R | R | 9 | 3 | | 27 | 0 |
| 27 | 70 | 25 | F | R | R | 57 | 52 | | 28 | 0 |
| 28 | 41 | 28 | M | R | R | 22 | 6 | | 26 | 0 |
| 29 | 47 | 28 | M | R | L | 59 | 54 | | 20 | 1 |
| 30 | 55 | 19 | M | R | L | 48 | 23 | | 24 | 2 |
| 31 | 76 | 20 | M | R | L | 52 | 53 | | 21 | 3 |
| 32 | 41 | 13 | M | R | L | 57 | 54 | | 23 | 0 |
| 33 | 63 | 29 | M | R | R | 31 | 19 | | 27 | 0 |
| 34 | 73 | 17 | M | R | R | 45 | 36 | | 25 | 0 |
| 35 | 62 | 11 | M | R | R | 49 | 50 | | 25 | 8 |

| **Late group**  Patients | age (years) | Time since stroke | gender | handedness | stroke hemisphere | FM-UE | ARAT | MoCa | GDS |
| --- | --- | --- | --- | --- | --- | --- | --- | --- | --- |
| 1 | 27 | 819 | F | R | R | 58 | 45 | 30 | 3 |
| 2 | 45 | 1649 | M | R | R | 13 | 3 | 30 | 0 |
| 3 | 68 | 1273 | M | R | R | 60 | 57 | 27 | 14 |
| 4 | 67 | 816 | M | R | R | 13 | 6 | 25 | 0 |
| 5 | 60 | 806 | M | R | L | 58 | 55 | 25 | 3 |
| 6 | 58 | 1178 | M | R | R | 48 | 44 | 25 | 2 |
| 7 | 48 | 734 | M | R | R | 12 | 3 | 29 | 0 |
| 8 | 65 | 2321 | M | R | R | 10 | 7 | 28 | 8 |
| 9 | 53 | 332 | F | L | R | 62 | 57 | 23 | 7 |
| 10 | 62 | 2680 | F | R | R | 18 | 3 | 25 | 1 |
| 11 | 58 | 739 | M | R | R | 30 | 6 | 20 | 1 |
| 12 | 78 | 253 | M | R | R | 50 | 43 | 20 | 4 |
| 13 | 50 | 1800 | M | R | R | 24 | 43 | 22 | 7 |
| 14 | 59 | 612 | M | R | R | 18 | 3 | 20 | 4 |
| 15 | 40 | 2191 | M | R | L | 61 | 55 | 27 | 3 |
| 16 | 55 | 1749 | F | R | L | 9 | 2 | 21 | 7 |
| 17 | 52 | 1077 | M | R | R | 65 | 56 | 26 | 4 |
| 18 | 68 | 212 | M | R | L | 65 | 57 | 28 | 0 |
| 19 | 49 | 221 | M | R | R | 27 | 14 | 29 | 8 |
| 20 | 52 | 251 | F | R | R | 36 | 14 | 28 | 1 |
| 21 | 57 | 506 | F | R | R | 15 | NaN | 26 | n.d. |
| 22 | 45 | 272 | F | n.d. | R | 33 | 25 | 28 | 14 |
| 23 | 38 | 216 | M | R | L | 61 | 57 | 26 | 0 |
| 24 | 58 | 2474 | F | R | R | 18 | 3 | 23 | 5 |
| 25 | 69 | 2900 | F | R | L | 64 | 57 | 27 | 1 |
| 26 | 39 | 3131 | F | R | L | 48 | 52 | 27 | 0 |
| 27 | 72 | 581 | M | R | R | 14 | 4 | 28 | 2 |
| 28 | 56 | 929 | M | R | R | 60 | 54 | 30 | 0 |
| 29 | 67 | 1088 | M | R | R | 22 | 24 | 29 | 5 |
| 30 | 64 | 10834 | M | R | L | 20 | 3 | 22 | 3 |

| **Controls** | age (years) | gender | handedness | MoCa | GDS |
| --- | --- | --- | --- | --- | --- |
| 1 | 31 | F | R | 29 | 0 |
| 2 | 59 | M | R | 29 | 0 |
| 3 | 79 | F | R | 29 | 0 |
| 4 | 79 | M | L | 26 | 0 |
| 5 | 61 | M | R | 27 | 0 |
| 6 | 61 | F | R | 26 | 0 |
| 7 | 42 | F | R | 27 | 0 |
| 8 | 62 | F | L | 30 | 0 |
| 9 | 28 | F | L | 29 | 0 |
| 10 | 63 | M | L | 28 | 0 |
| 11 | 47 | F | L | 29 | 0 |
| 12 | 52 | F | L | NaN | 0 |
| 13 | 48 | F | L | NaN | 0 |
| 14 | 65 | F | L | 29 | 0 |
| 15 | 60 | F | L | NaN | 0 |
| 16 | 52 | F | R | NaN | 1 |
| 17 | 64 | F | L | 24 | 0 |
