## Supplementary material for "Reinforcement Learning Is Impaired in the Sub-acute Post-stroke Period": Table 3

|  | **Reinforcement Task** | | | **Error-based Task** | | |
| --- | --- | --- | --- | --- | --- | --- |
| Group | Early | Late | Control | Early | Late | Control |
| Total Learning | -4.9° ±9.4 | -7.5° ±9.1 | -11.1° ±8.3 | -10.82° ±4.9 | -12.1° ±7.2 | -13° ±1.9 |
| End Perturbation | -5.7° ±11.1 | -9.3° ±10.3 | -13.1° ±9.3 | -12.54° ±2.1 | -13.44° ±3.9 | -13.1° ±3.5 |
| Early Retention | 1.1° ±5.5 | 1.9° ±7.0 | 3.4° ±7.8 | 4.5° ±8.8 | 6.1° ±8.6 | 7.2° ±6.4 |
| Subgroup |  |  | **-** |  |  | **-** |
| End Perturbation | -2.7° ±9.4 | -9.6° ±7.0 | **-** | -12.11° ±3.6 | -12.87° ±4.8 | **-** |
| Early Retention | 0.8° ±4.3 | 1.9° ±5.5 | **-** | 5.2° ±7.7 | 9.2° ±5.4 | **-** |

**Table 3 Outcomes per task of each group**

Median ±indicates standard deviation across participants. *Total Learning*: average reaching angle for the last 40 trials of perturbation minus the first 40 trials of the baseline, *End Perturbation*: average reaching angle for the last 40 trials of perturbation, *Early Retention:* first 40 trials of the retention block. *Subgroup analysis:* participants from both stroke groups with matched *Baseline* performance.
